# Nephronectin-Integrin α8 signaling is required for proper migration of periocular neural crest cells during chick corneal development

**DOI:** 10.1101/2021.10.13.464255

**Authors:** Justin Ma, Lian Bi, James Spurlin, Peter Lwigale

## Abstract

During development, cells aggregate at tissue boundaries to form normal tissue architecture of organs. However, how cells are segregated into tissue precursors remains largely unknown. Cornea development is a perfect example of this process whereby neural crest cells aggregate in the periocular region prior to their migration and differentiation into corneal cells. Our recent RNA-Seq analysis identified upregulation of Nephronectin (Npnt) transcripts during early stages of corneal development where its function has not been investigated. We found that Npnt mRNA and protein are expressed by various ocular tissues including the migratory periocular neural crest (pNC), which also express the integrin alpha 8 (Itgα8) receptor. Knockdown of either *Npnt* or *Itgα8* attenuated cornea development, whereas overexpression of *Npnt* resulted in cornea thickening. Moreover, overexpression of Npnt variants lacking RGD binding sites did not affect corneal thickness. Neither the knockdown or augmentation of Npnt caused significant changes in cell proliferation, suggesting that Npnt directs pNC migration into the cornea. *In vitro* analyses showed that Npnt promotes pNC migration from explanted periocular mesenchyme, which requires Itgα8. Combined, these findings show that Npnt specifies and tunes cell migration into the presumptive cornea ECM by providing a substrate for Itgα8-positive pNC cells.

## INTRODUCTION

In vertebrates, the cornea is comprised of three cellular layers: the epithelium, stroma and endothelium. The corneal epithelium is derived from the ectoderm, whereas the stromal keratocytes and corneal endothelium are derived from cranial neural crest cells (Lwigale et al., 2005). In birds and humans, neural crest migration from the periocular region into the presumptive cornea occurs in two waves (Feneck et al., 2020; E. D. Hay, 1980). The first wave forms a monolayer of the nascent corneal endothelium, which is accompanied by a second wave of mesenchymal migration into the acellular matrix between endothelium and ectoderm to form the cornea stroma (Hay and Revel, 1969; Noden, 1978; Lwigale et al., 2005; Creuzet et al., 2005; Feneck et al., 2020). Several mechanisms pertaining to the role of growth factors and guidance cues have been identified (Beebe and Coats, 2000; Saika et al., 2001; Lwigale and Bronner-Fraser, 2009; Choi et al., 2014), but the functional role of ECM/integrin signaling in establishing the cornea remains unclear.

Among the initial effects of the signaling that takes place between the nascent ocular tissues is the synthesis of the corneal extracellular matrix (ECM) by the corneal epithelium in response to inductive signals from the lens vesicle to form the primary stroma (Hay and Revel, 1969; Hendrix et al., 1982; Fitch et al., 1988). In birds and humans, the primary stroma serves an important role as a scaffold for pNC migration throughout the process of corneal development (Hay and Revel, 1969; Bard and Hay, 1975; Quantock and Young, 2008). It consists of multiple ECM proteins including hyaluronan (Toole and Trelstad, 1971), collagen type I and II (Hayashi et al., 1988; Fitch et al., 1988), laminin (Doane et al., 1996), and fibronectin (Fn) (Fitch et al., 1991). Following the second wave of pNC migration into the developing cornea, the primary stroma is concomitantly replaced by the secondary stroma that is synthesized by differentiating stromal keratocytes. The secondary stroma comprises the bulk of the adult cornea and consists of collagens and proteoglycans (Hayashi et al., 1988; Quantock and Young, 2008) that are arranged in patterns that result in transparency (Chen et al., 2015). Over the past decades, the role of collagens and proteoglycans have been the focus of several investigations due to their indispensable functions in corneal transparency. However, most of these studies were conducted in rodents, in which early corneal development does not involve the primary stroma (Pei and Rhodin, 1970; Haustein, 1983; Feneck et al., 2019). Although the ECM plays a critical role during organogenesis by providing cell adhesion substrate, sequestering signaling molecules, providing structural support and mechanical cues (Hynes, 2014), the function of the primary stroma during early corneal development remains to be elucidated.

Our recent RNASeq analysis of pNC differentiation into corneal cells identified novel expression and upregulation of Nephronectin (Npnt) transcripts during corneal development. Npnt is a 70-90 kDa protein that was discovered as a novel ECM ligand for integrin α8β1 (Itgα8β1) during kidney development (Brandenberger et al., 2001), and at the same time, as a protein secreted by preosteoblast cells (Morimura et al., 2001). It consists of five epidermal growth factor (EGF)-like domains in the N-terminal, a central region containing an Arg-Gly-Asp (RGD) sequence, and a meprin-A5 protein-receptor protein tyrosine phosphatase μ (MAM) domain in the C-terminal. The EGF-like and RGD domains have been functionally characterized and shown to play critical roles in development and tissue homeostasis. The RGD domain signals through α8β1 receptor during epithelial-mesenchymal interactions involved in kidney development and maintenance (Miner, 2001, Linton et al., 2007; Müller et al., 1997, Sato 2009, Cheng et al., 2008, Inagi et al., 2017, Müller-Deile et al., 2017, Zimmerman et al., 2018). These observations in mice were recently confirmed by the identification of recessive mutation in the Npnt gene that caused bilateral kidney agenesis in human fetuses (Dai et al., 2021). The EGF-like domains of Npnt are associated with inducing differentiation and proliferation in osteoblasts and dental stem cells through the activation of mitogen-activated protein kinase (MAPK) pathways (Fang et al., 2010; Kahai et al., 2010; Arai et al., 2017), and to induce vascular endothelial cell migration via phosphorylation of extracellular signal-regulated kinases (ERK) and p38MAPK (Kuek et al., 2016). Recent studies have also implicated Npnt to play a role in heart development (Patra et al., 2011), attachment of the arrector pili muscle to hair follicles (Fujiwara et al., 2011), and during forelimb formation in amphibians (Abu-Daya et al., 2011). Other studies have also demonstrated that Npnt plays a potential role in diseases such as chronic obstructive pulmonary disease (COPD) (Saferali et al., 2020), diabetic glomerulosclerosis (Nakatani et al., 2012), Fraser syndrome (Kiyozumi et al., 2012), and in various cancers (Magnussen et al., 2021). Despite the pleiotropic functions of Npnt in development and disease, its expression and function have not yet been described in the cornea.

Inspired by our observation that Npnt transcripts were upregulated during corneal development, we sought to establish the role of Npnt during avian corneal development. We first characterized the spatiotemporal expression of Npnt mRNA and protein at stages that correspond with pNC migration. Next, we used RCAS-mediated gene knockdown to investigate the role of *Npnt*. We identified Itgα8 as a potential receptor for Npnt during cornea development and confirmed its role in pNC using *in vitro* migration assay coupled with peptide inhibition, and *in vivo* using gene knockdown. Lastly, we performed misexpression studies using the either the full-length, RGD mutant, and truncated versions of Npnt to further confirm its role and the signaling mechanism during corneal development. Together, our findings show that Npnt secreted into the ECM of the nascent cornea provides a substrate that promotes migration of Itgα8-positive cells, thus segregating the cornea progenitors from other pNC cells.

## RESULTS

### Npnt mRNA and Protein are expressed during early ocular development

Our previous RNA-seq analysis of gene expression during pNC differentiation into corneal cells identified upregulation of *Npnt* transcripts during chick corneal development (Bi and Lwigale, 2019). To define the expression of Npnt mRNA and protein distribution during this process, we performed section *in situ* hybridization and immunohistology, respectively. Consistent with our RNA-seq data (Bi & Lwigale, 2019), we found that *Npnt* is undetectable in the pNC by embryonic day (E) 4 and in the presumptive corneal endothelium at E5, but it is primarily expressed in the retinal pigment epithelium (RPE) and presumptive lens fiber cells at these time points (Fig. 1A, B). However, by E6, we found that in addition to the RPE and lens, *Npnt* is vividly expressed by the second wave of migratory pNC that eventually differentiate into the stromal keratocytes (Fig. 1C, arrowhead). Expression of *Npnt* persisted in the corneal stroma through E12 (Fig. S1A-C), but it was undetectable at E15 (data not shown).

**Fig. 1.**
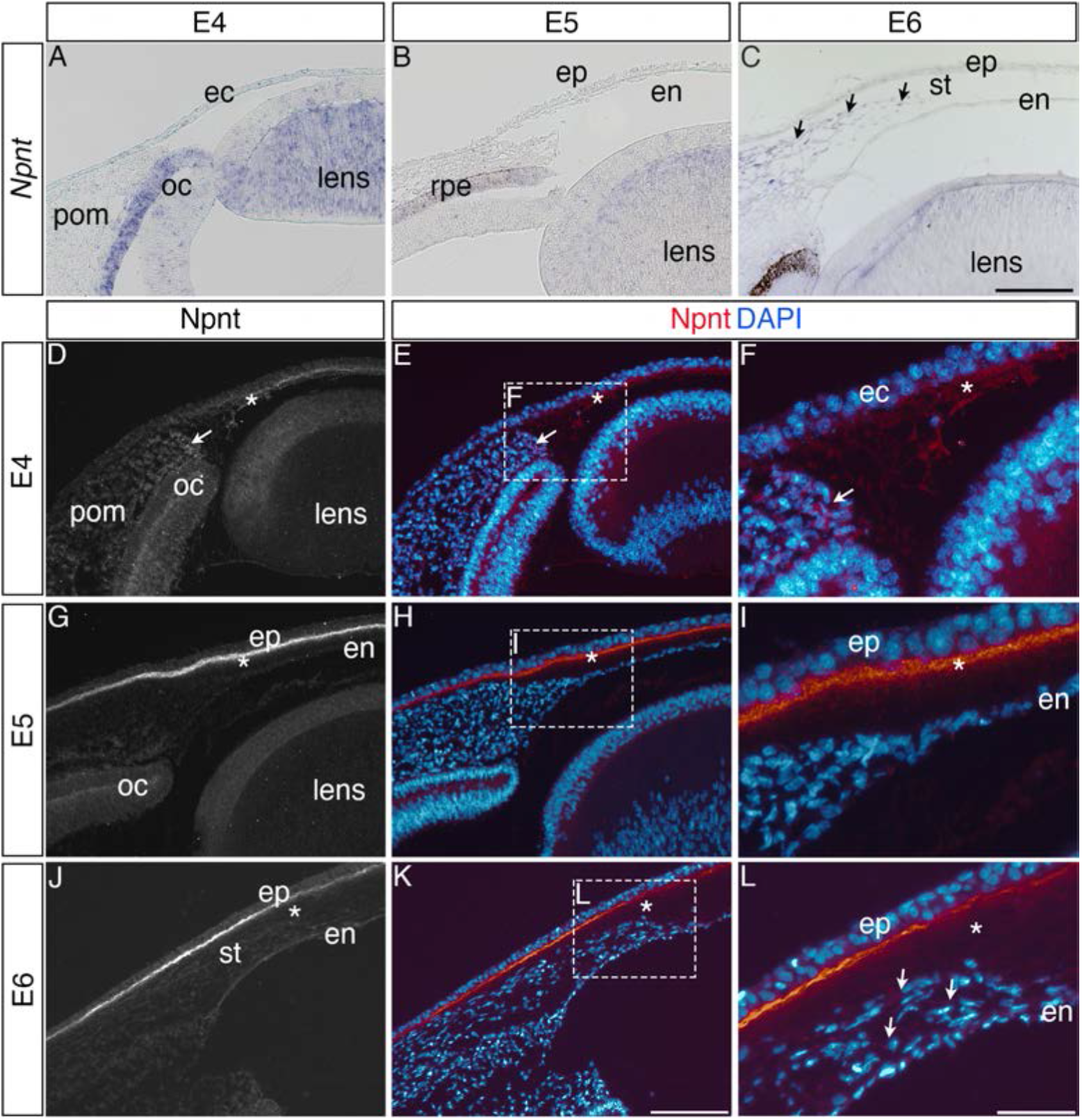
Npnt expression during early ocular development. Expression of Npnt mRNA and protein were examined via section in situ hybridization (A-C) or immunohistochemistry (D-L). (A-B) *Npnt* expression in the retina pigment epithelium layer of the optic cup and region of the presumptive lens fiber cells at E4 and E5. (C) Initial expression of *Npnt* by pNC is observed during the second wave of migration into the stroma (black arrows). (D-F) At E4, Npnt protein was detected in the optic cup, lens epithelium, in the periocular mesenchyme proximal to the presumptive cornea region (arrow), and in the matrix of the primary stroma (asterisk). (G-I) At E5, vivid expression of Npnt protein is localized in the primary stroma adjacent to the corneal epithelium and diffusely throughout the primary stroma (asterisk), and persists at low levels in the optic cup and lens epithelium. At E6, vivid expression of Npnt protein persists in the primary stroma adjacent to the corneal epithelium and it remains diffusely expressed throughout the primary stroma (asterisk). At this time, low expression of Npnt protein is also observed in the migratory periocular neural crest cells invading the primary stroma (arrows in L). Abbreviations: ec, ectoderm; oc, optic cup; pom, periocular mesenchyme; rpe, retinal pigment epithelium; st, stroma; en, corneal endothelium; ep, corneal epithelium. Scale bars: 100 μm.

Because Npnt is a secreted protein that plays a role in cell migration (Kuek et al., 2016; Magnussen et al., 2021; Yamada & Kamijo, 2016), we examined its localization at similar time points during cornea development. At E4, Npnt is localized in the optic cup, lens epithelium, surrounding the pNC cells adjacent to the nascent cornea (Fig. 1D-F, arrow), and in the primary stroma of the cornea (Fig. 1D-F, asterisk). At E5, strong Npnt staining is observed in the primary stroma (Fig. 1G-I, asterisk). By E6, Npnt is localized to the basement membrane of the corneal epithelium and diffusely throughout the primary stroma (Fig. 1J-L). At this time, the migratory pNC also stain positive for Npnt (Fig 1L, arrowheads). Strong expression of Npnt persists in the basement membrane of the corneal epithelium through E9 (Fig. S1 C,D), and it is faintly detectable in the corneal epithelial cells by E12 (Fig. S1 E). These data show that expression of Npnt coincides with pNC ingression into the cornea, implicating a potential role during development.

### Knockdown of *Npnt* disrupts corneal development

To investigate the function of Npnt during early corneal development, we generated several RCAS-GFP-Npnt-shRNA (*Npnt^kd^*) constructs in chick DF-1 cells (Supplementary Fig. S2). The construct with the highest knockdown of *Npnt* was used for the experiments. Constructs containing only GFP (RCAS-GFP) or scrambled shRNA (RCAS-GFP-Scr-shRNA) were used as control. The viral constructs were injected to cover the entire cranial region of HH7-8 (Hamburger & Hamilton, 1951) chick embryos (Fig. 2A). Using this approach, both the neural progenitors of the optic cup and neural crest cells, as well as the ectodermal progenitors of the lens, where *Npnt* is expressed (Fig. 1A-C), are infected by the viral constructs. Embryos were re-incubated and collected at E7, then the anterior eyes were screened for GFP as an indicator for the extent of viral infection. Only the eyes that showed robust GFP expression were evaluated for knockdown of *Npnt* expression (Fig. 2A). Histological analysis of E7 corneas based on Hematoxylin and Eosin staining revealed that knockdown of *Npnt* (N=4) resulted in reduction of corneal thickness (Fig. 2C) compared to control (N=5; Fig. 2B). Measurements taken in the mid-corneal regions (Fig. 2D, 2E; arrowheads) showed significant reduction in corneal thickness in *Npnt^kd^* corneas (Fig. 2F). These data suggest that Npnt plays a critical role during normal formation of the cornea.

**Fig. 2.**
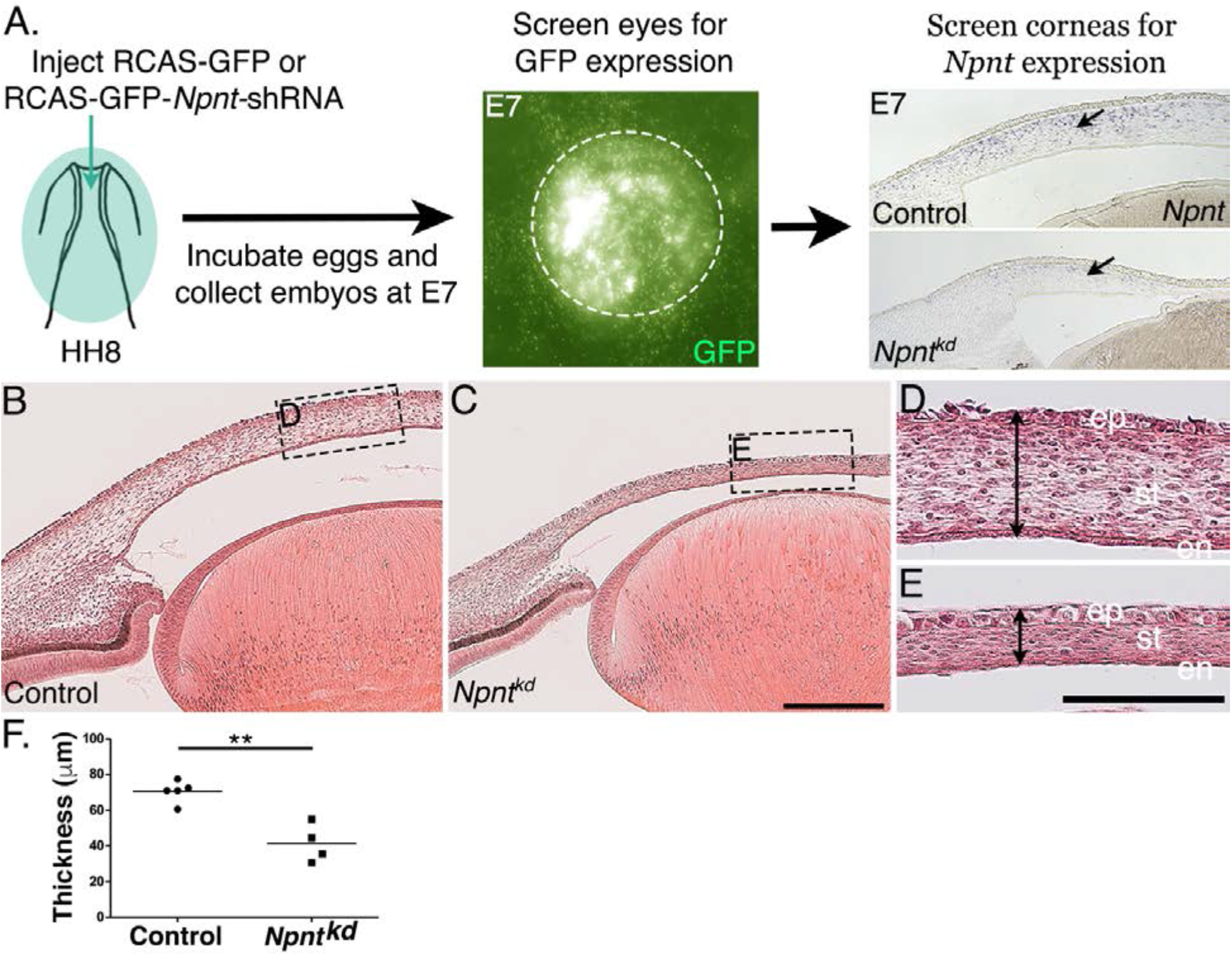
Corneal thickness is reduced in *Npnt^kd^* corneas. (A) Schematic of *in ovo* injection of viral constructs (green) to cover the anterior region of stage 8 chick embryo. Following 7 days of incubation, embryos were screened for GFP expression in the anterior eye region. Knockdown was verified by section in situ hybridization, which revealed reduced expression of *Npnt* in *Npnt^kd^* cornea compared with control. (B-E) Hematoxylin and Eosin staining showing control (B,D) and thinner *Npnt^kd^* corneas (C,E). Statistical analysis on measurements taken from (N=5 control and N=4 *Npnt^kd^* corneas) revealed: (F) Significant reduction in thickness of *Npnt^kd^* corneas, * * indicates p < 0.01. Abbreviations: ep, corneal epithelium; st, stroma; en, corneal endothelium;. Scale bars: 100 μm (B,C), 100 μm (D,E).

### Itgα8 is the receptor for Npnt during cornea development and plays a role in pNC migration

Next, we reasoned that because knockdown of *Npnt* causes a significant reduction in corneal thickness, one or more of its receptors should be expressed by the migratory pNC during cornea development. In this study, we focused on integrin α8β1 because of its strong affinity for Npnt (Brandenberger et al., 2001). Although integrin α8β1 was observed to promote spreading of trunk neural crest *in vitro* (Testaz et al., 1999), little is known about its expression and function in the cranial neural crest. Our analysis by *in situ* hybridization revealed that *Itgα8* was expressed in the leading edge of the periocular mesenchyme prior to pNC migration into the primary stroma of the nascent cornea (Fig. 3A, arrow). Subsequently, *Itgα8* was maintained in the periocular mesenchyme but also expressed in the corneal endothelium at E5 (Fig. 3B, arrows) and stroma at E6 (Fig. 3C).

**Fig. 3.**
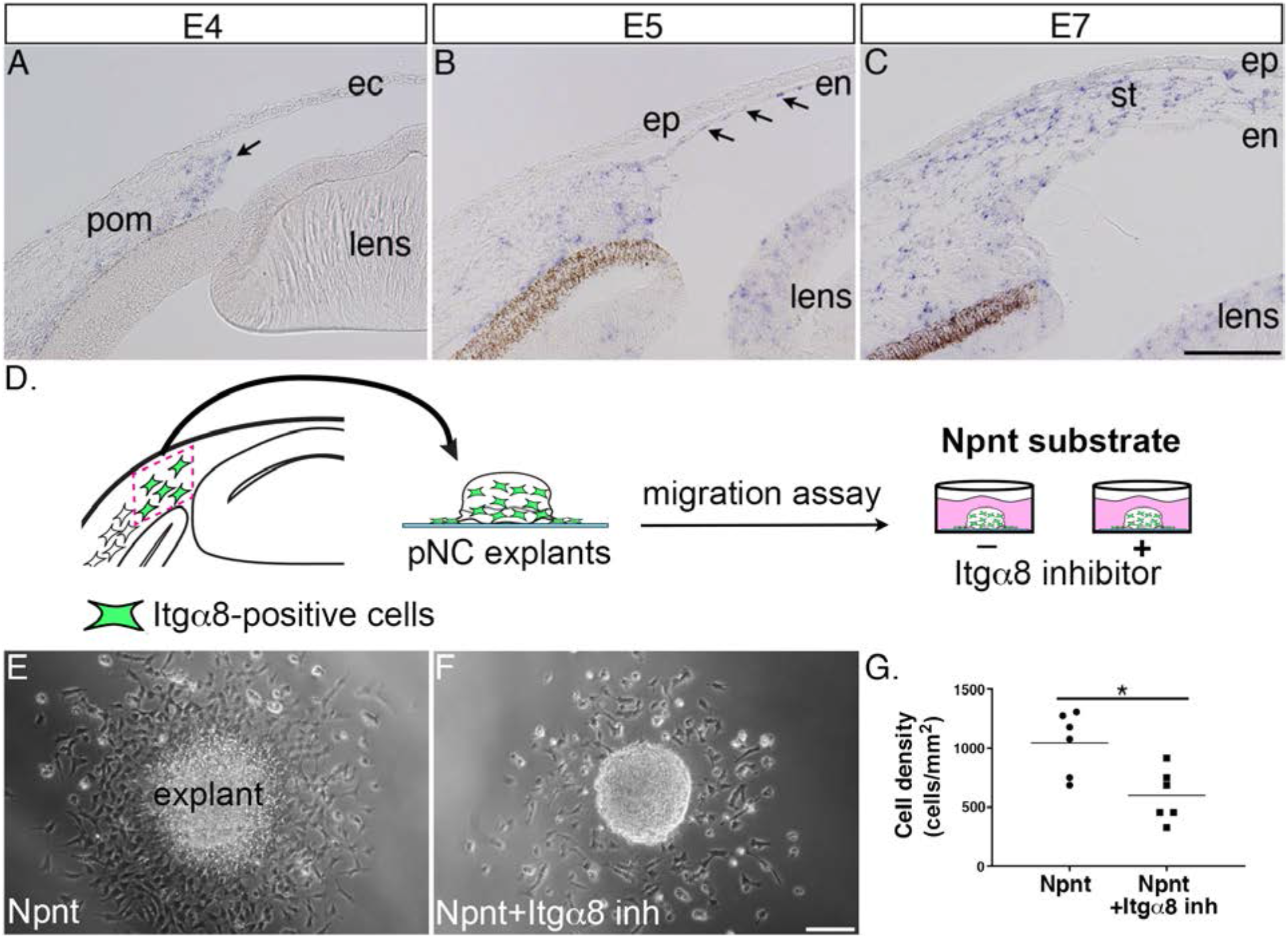
Itgα8 is expressed pNC during cornea development and it plays a role in cell migration. (A) Expression of *Itgα8* is observed in pNC prior to their migration into the cornea at E4 (arrow). (B) *Itgα8* is subsequently expressed in the corneal endothelium at E5 (arrows), and (C) by the migratory pNC in the corneal stroma at E6. (D) Schematic showing the isolation of periocular mesenchyme used for generating pNC explants for *in vitro* migration on Npnt-coated substrate in the presence or absence of Itgα8 inhibitor. (E) Explant cultured on Npnt substrate showing robust cell migration after 12 hours. (F) Explant cultured on Npnt substrate in the presence of Itgα8 inhibitor showing fewer cell migration after 12 hours. (G) Statistical analysis performed on (N=6 explants on Npnt substrate) and (N=6 explants on Npnt substrate+Inhibitor) revealed significant reduction in cell density of migratory cells in the presence of the inhibitor. * indicates p < 0.05. Abbreviations: ec, ectoderm; pom, periocular mesenchyme; en, corneal endothelium; ep, corneal epithelium; st, stroma. Scale bars: 100 μm.

Given the striking expression of *Itgα8* by the migratory pNC, we first assessed the potential for Itgα8-Npnt signaling *in vitro* using explanted periocular mesenchyme from the leading edge adjacent to the presumptive cornea (Fig. 3A and 3D). Mesenchyme explants were cultured on slides coated with Npnt in the presence or absence of a peptide inhibitor previously shown to specifically inhibit binding of α8β1 with Npnt (Fig. 3D) (Sato et al., 2009). In the absence of the inhibitor, explants attached to the Npnt-coated slides and numerous migratory cells formed a halo around the explant within 12-17 hours of incubation (Fig. 3E; Movie 1). In contrast, we observed that treatment with the Itgα8 inhibitor significantly decreased the density of migratory cells from the explant (Fig. 3F, 3G; Movie 2). These results indicate that α8β1 functions as a receptor for Npnt signaling in the presumptive corneal pNC, and that it is required for their migration.

### Knockdown of *Itgα8* disrupts corneal development

Given that *Itgα8*-expressing pNC appear to be in direct contact with Npnt secreted into the primary stroma, we wanted to characterize the role of Itgα8 during corneal development. For this analysis, we generated and tested several RCAS-GFP-Itgα8-shRNA constructs and validated one that showed highest knockdown of *Itgα8* in DF-1 cells and *in vivo* (Supplementary Fig. S2 and Fig. S3). As a first step, we analyzed the effect of *Itgα8* knockdown on the first wave of pNC migration that forms the corneal endothelium at E5. We found that at this timepoint relatively fewer GFP expressing cells appeared to migrate into the *Itgα8^kd^* cornea (Fig. 4B, arrows) compared to RCAS-GFP control, which showed robust occupation of GFP-positive cells in the corneal endothelium (Fig. 4A). Histological analysis at E7 revealed reduction in corneal thickness following *Itgα8* knockdown (Fig. 4D) compared to control corneas (Fig. 4C). Measurements and quantification of cells in the mid-corneal regions showed an overall significant reduction in corneal thickness and cell number, but no difference in cell density between *Itgα8^kd^* and control corneas (Fig. 4E). Furthermore, we performed a BrdU assay and observed no significant differences in cell proliferation between *Itgα8^kd^* (Fig. 4G) and control corneas (Fig. 4F). These results indicate that Itgα8 is required for pNC migration into the cornea. Given that the corneal defects following knockdown of *Itgα8* correlate with *Npnt* knockdown, our results suggest that Npnt-Itgα8 signaling plays an important role during pNC migration into the cornea.

**Fig. 4.**
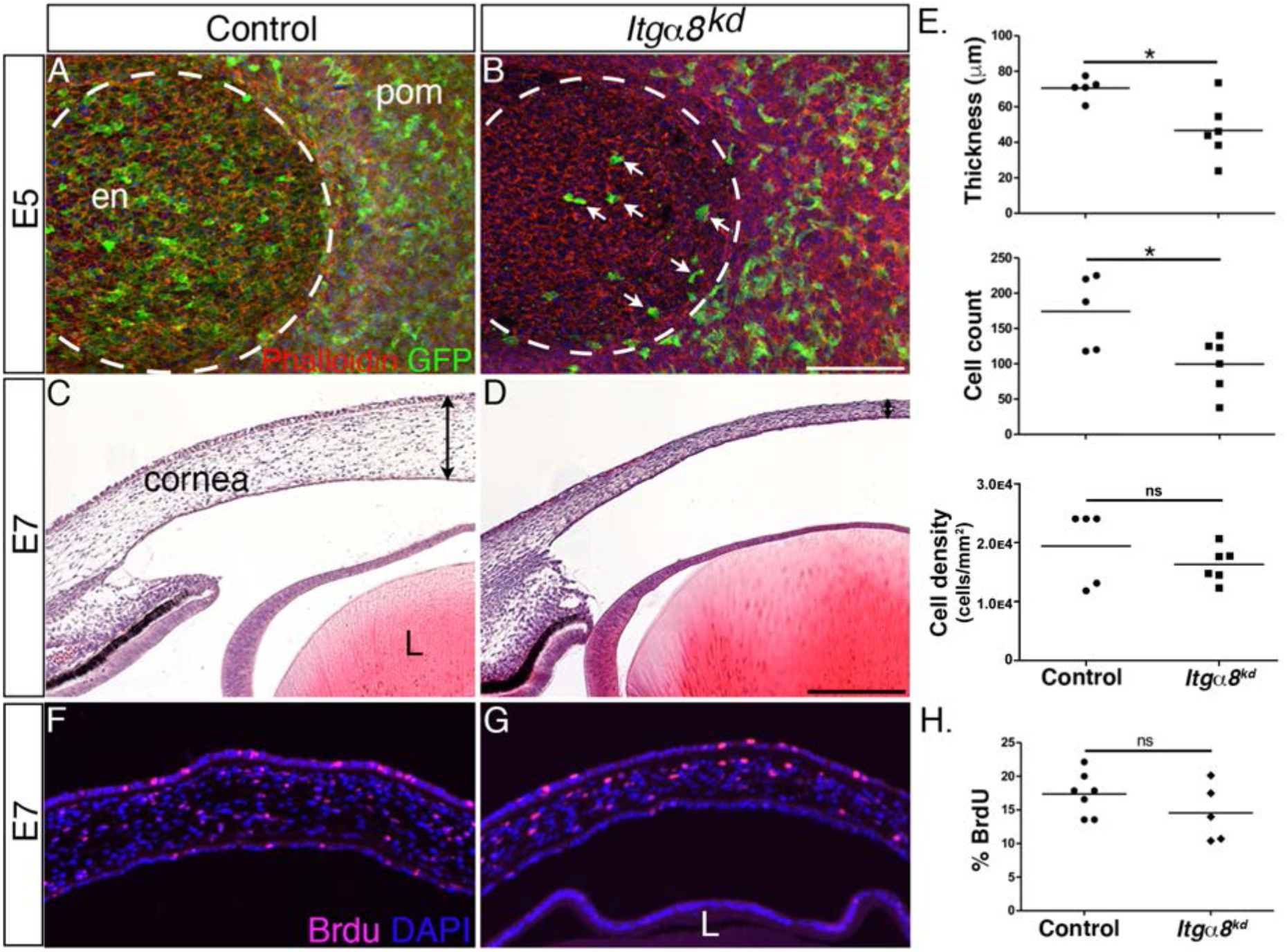
Knockdown of Itgα8 reduces pNC migration and results in reduced corneal thickness. (A-B) Whole-mount E5 anterior eyes immunostained for GFP and counterstained with phalloidin to reveal cell membranes. (A) Control eye showing robust expression of GFP by pNC cells that migrated into the corneal region to form the endothelial layer. (B) *Itgα8^kd^* eye showing that relatively fewer GFP cells migrated into the cornea (arrows). Dotted lines demark the boundary between the cornea and periocular mesenchyme. (C-D) Hematoxylin and Eosin staining of E7 corneal section showing (C) normal corneal thickness in control and (D) reduced corneal thickness in *Itgα*8^kd^ embryos. Double-sided arrows indicate corneal thickness. (E) Statistical analysis of measurements taken from (N=5 control and N=6 *Itgα8^kd^* corneas) revealed significant reduction in thickness and cell count, and no difference in cell density in *Itgα*8^kd^ corneas, * indicates p < 0.05. (F-G) BrdU immunofluorescent analysis of cell proliferation in E7 corneal sections. Quantification of BrdU-positive cells in the corneal stroma was performed by normalizing to the total number of DAPI-positive cells. (H) Statistical analysis from N=7 control and N=5 *Itgα8^kd^* revealed a no difference between control and *Itgα8*^kd^ corneas. ns, not significant. Abbreviations: en, corneal endothelium; pom, periocular mesenchyme; L, lens. Scale bars: 100 μm.

### Overexpression of *Npnt* causes corneal thickening

Given that knockdown of both *Npnt* and *Itgα8* resulted in corneal thinning, combined with our observation that *Npnt* is expressed during the second wave of migration at E6, we investigated whether overexpression of the *Npnt* affects cornea development. We generated and tested viral constructs containing the full-length Npnt gene (*Npnt^oe^*) in DF1 cells (Fig. S2). Control and *Npnt^oe^* constructs were injected in HH7-8 embryos as described (Fig. 2), and the corneas were collected for analysis at E7, E9, and E15. First, we confirmed the overexpression of Npnt mRNA and protein in the corneas. At E9, expression of *Npnt* was localized in the corneal stroma during normal development (Fig. 5A). However, *Npnt^oe^* corneas showed robust expression of *Npnt* in the stroma, as well as ectopic expression in the corneal epithelium and endothelium, and the lens (Fig. 5D). At this stage, strong staining for Npnt protein was only observed in the basement membrane of the corneal epithelium and low diffused staining in the stroma of control corneas (Fig. 5B). In contrast, the *Npnt^oe^* corneas showed strong ectopic staining for Npnt in the corneal epithelium and endothelium, and a substantial increase in protein expression in the stroma compared to the control (Fig. 5E).

**Fig. 5.**
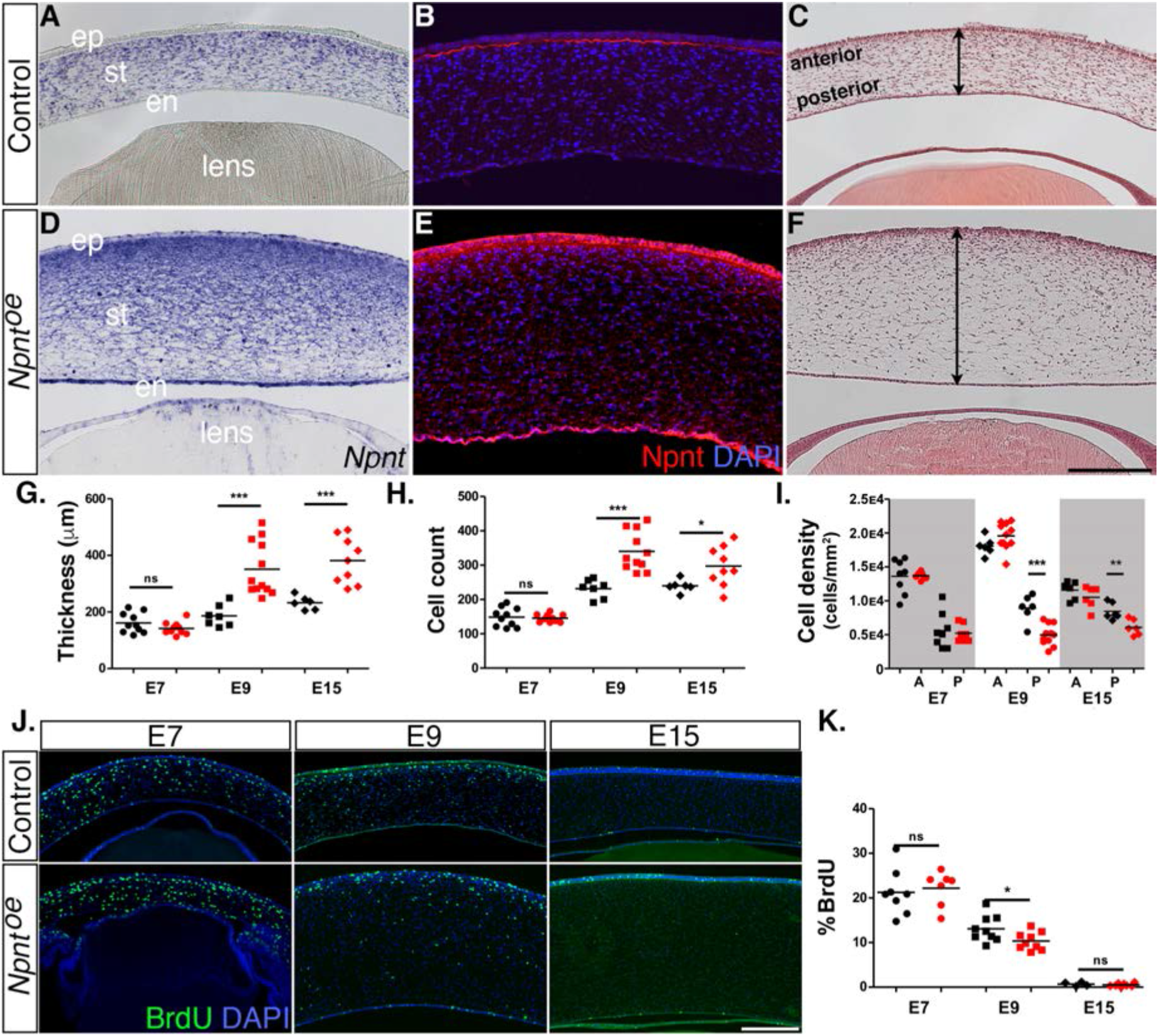
Effects of overexpression of Npnt during corneal development. Embryos were injected with RCAS virus expressing GFP alone (control) or GFP and the full-length Npnt protein, and corneas were analyzed at the developmental stages indicated. (A-F) Representative corneal sections from E9 control (A-C) and *Npnt^oe^* corneas (D-F) showing levels of Npnt transcript (A,D) and protein (B,E) expression, and H&E staining indicating corneal thickness (double sided arrows, C,F). (G,H) Measurements for corneal thickness and cell counts were taken from: E7, N=10 control, N=10 *Npnt^oe^*; E9; N=7 control, N=12 *Npnt^oe^*; E15; N=6 control, N=9 *Npnt^oe^*. Bar graphs show no difference at E7, but a significant increase at E9 and E15 in corneal thickness (G), and corneal cells (H). (I) Cell densities were determined from: E7, N=8 control, N=7 *Npnt^oe^*; E9; N=6 control, N=11 *Npnt^oe^*; E15; N=7 control, N=6 *Npnt^oe^*. Bar graph shows that there were no differences at E7 and anterior cell densities at E9 and E15, but the posterior cell densities were significantly decreased. (J and K) BrdU analysis and quantification of cell proliferation in corneal sections taken from E7, N=8 control, N=7 *Npnt^oe^*; E9; N=9 control, N=9 *Npnt^oe^*; E15; N=7 control, N=6 *Npnt^oe^*. No significant differences were observed at E7 and E15, but there was a significant reduction at E9. ns, not significant; * indicates p < 0.05; ** indicates p < 0.01; and *** indicates p < 0.001. Abbreviations: ep, corneal epithelium; st, stroma; en, corneal endothelium. Scale bars: 100 μm.

Next, we performed histological analysis on E7, E9, and E15 corneas to determine whether there were any morphological differences between the control and *Npnt^oe^* corneas. Our analysis did not reveal any significant differences in corneal thickness and cell number at E7 (Fig. 5G, H). By contrast, cross-sections of E9 *Npnt^oe^* corneas revealed significant thickening and increase in the cell number (Fig. 5F, G, H) compared with control (Fig. 5C, G, H). Similar increases in corneal thickness and cell number were observed at E15 (Fig. 5G, H). Given that the keratocyte density is higher in the anterior stroma than the posterior stroma (Patel et al., 2001; Berlau et al., 2002), we quantified the cell densities in these regions. Overall, we observed relatively higher cell densities in the anterior stroma of E7, E9, and E15 corneas and there were no significant differences between *Npnt^oe^* and control corneas (Fig. 5I). Posterior cell density was not affected in E7 *Npnt^oe^* corneas, but there was significant reduction at E9 and E15.

To determine whether elevated cell proliferation was a contributing factor to the increased corneal thickness in *Npnt^oe^*, we examined corneas at E7, E9, and E15 by performing BrdU labeling and immunofluorescent detection. Overall, we observed cell proliferation in all corneal layers, and there was a decreasing trend in the percentage of labeled cells in the stroma as development progressed from E7 to E15 (Fig. 5J, K). However, our results revealed no significant increase in cell proliferation at E7 and E15. Surprisingly, there was a significant reduction in cell proliferation in E9 *Npnt^oe^* corneas despite their increase in thickness (Fig. 5K). Combined, our data indicate that overexpression of *Npnt* increases transcript and protein expression in the cornea, which results in increased cell number in the stroma and corneal thickness. Since these increases were not correlated with increased cell proliferation, our data suggest that the changes in corneal thickness are due to augmented pNC migration into the cornea caused by excessive expression of Npnt.

### The RGD domain mediates Npnt signaling in pNC during corneal development

Npnt mediates signal transduction in development and cancer by binding to various receptors through the EGF-like and RGD domains (Arai et al., 2017; Kahai et al., 2010; Kuek et al., 2016; Linton et al., 2007). The marked reductions in corneal thickness observed following knockdown of both *Npnt* and *Itgα8* (Fig.2 and Fig.3) suggest that Npnt functions via the RGD domain during pNC migration. Given that overexpression of the full-length Npnt protein caused corneal thickening, we generated an Npnt-RAE version of the protein in which the RGD domain was mutated by substitution with RAE sequence, and an Npnt-EGF version in which both the RGD and MAM domains were truncated (Fig. 6A). Constructs were injected in HH7-8 embryos and E9 corneas were collected, sectioned, and analyzed following DAPI staining. Contrary to our observations following overexpression of the full-length protein (Fig.5), neither the Npnt-RAE or Npnt-EGF versions caused a significant change in corneal thickness compared with the control (Fig. 6B, C). Interestingly, there was a significant reduction in cell count and density in Npnt-RAE corneas although no difference in these values was observed for Npnt-EGF (Fig. 6D, E). The absence of corneal thickening following overexpression of either Npnt-RAE or Npnt-EGF constructs further suggest that the RGD domain plays an essential role in this process. Our results also indicate that the EGF-like domain does not play an essential role during for pNC migration into the cornea.

**Fig. 6.**
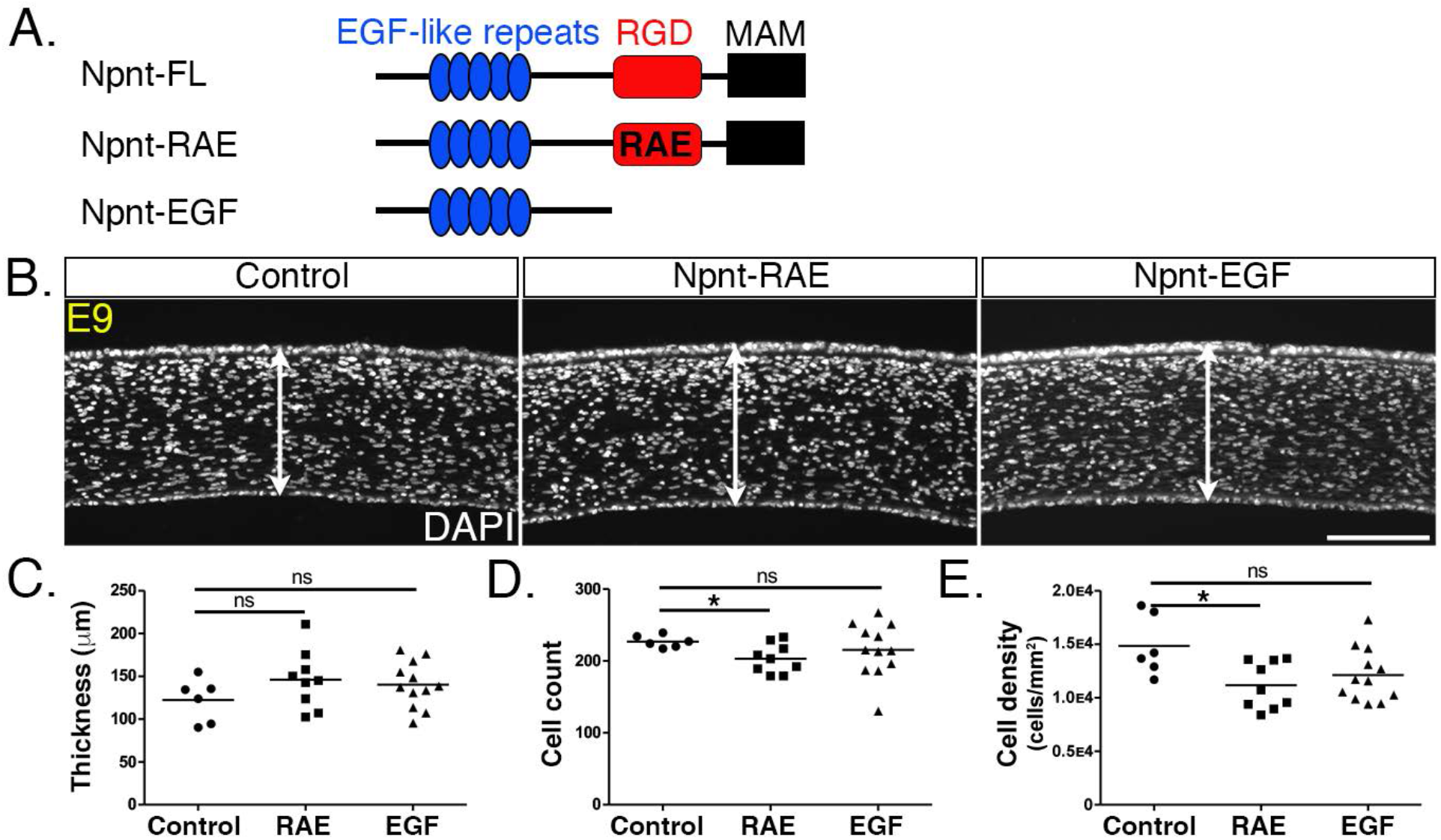
Overexpression of versions of Npnt with either mutant or truncated RGD domains does not increase corneal thickness. (A) Schematic showing the full-length Npnt, Npnt with mutated RGD to RAE sequence, and the truncated version containing only the EGF domain. (B) representative sections of E9 corneas showing corneal thicknesses (double sided arrows) following overexpression of control, RGD mutant, and truncated versions of Npnt. (C-E) Quantification of measurements taken from N=5 control, N=9 RAE, and N=12 EGF showing: (C) no significant differences in corneal thickness, (D,E) significant reduction in cell count and density in RGD mutant, but no difference in the truncated version. ns, not significant; * indicates p < 0.05. Scale bar: 100 μm.

## DISCUSSION

Very little is known about the function of the corneal ECM during early development. Here, we focus on Npnt, which we recently found in our RNA-seq study (Bi & Lwigale, 2019) to be upregulated during early development of the chick cornea. Specifically, we have identified novel expression of Npnt and its Itgα8 receptor at critical timepoints of corneal development. Our data reveal that disruption of Npnt/Itgα8 signaling either *in ovo* or *in vitro* perturbs pNC migration that subsequently result in corneal thickness defects, but formation of the cellular layers is not affected. Our model (Fig. 7) provides a context in which Npnt functions in the presence of Fn during corneal development. Fn is a well-known substrate for neural crest cell migration (Alfandari et al., 2003; Bronner-Fraser, 1986; Newgreen & Thiery, 1980) that is robustly expressed in the periocular mesenchyme and cornea (Doane et al., 1996; Kurkinen et al., 1979). In comparison, Npnt is expressed in an increasing gradient from the edge of the periocular region towards the cornea (Fig. 7A). We posit that all pNC cells express α5β1 and previously showed that they robustly migrate on Fn substrate *in vitro* (Lwigale & Bronner-Fraser, 2009). However, *in vivo*, it is most likely that the pNC which express both α5β1 and α8β1, and thus can respond to Npnt and Fn, migrate into the cornea. Based on our current study, we conclude that Npnt/Itgα8 signaling plays an essential role in pNC migration during corneal development.

**Fig. 7.**
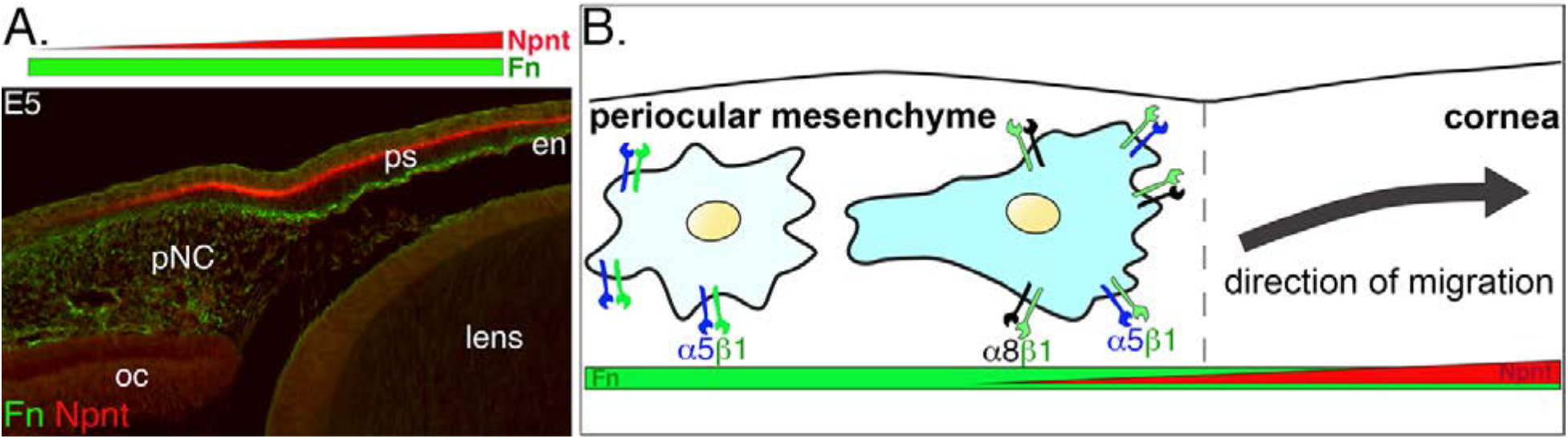
Dynamics of pNC response to the ECM in context of the expression of Npnt and Fn during corneal development. (A) Cross-section of E5 eye immunostained for Npnt and Fn. Npnt appears in an increasing gradient from the periocular region into the cornea, whereas Fn stains both the periocular mesenchyme and cornea. (B) All pNC respond to Fn via expression of α5β1, but a subpopulation of pNC that reside in the region adjacent to the presumptive cornea express both α5β1 and α8β1, and become competent to also read the additional gradient of Npnt in the ECM, thus migrating into the corneal region. Abbreviations: pNC, periocular neural crest; oc, optic cup; ps, primary stroma; en, corneal endothelium.

### Npnt and Itgα8 in the cornea

While Npnt is associated with kidney, teeth, and bone development (Arai et al., 2017; Kuek et al., 2016; Linton et al., 2007), only one reference was made to its expression in the mouse lens during ocular development (Brandenberger et al., 2001). First, we confirmed that *Npnt* is expressed in the migratory pNC and showed that it is also localized in the optic cup and lens vesicle. Our immunohistochemistry analysis revealed that Npnt protein staining consistently corresponded with the mRNA expression during corneal development. In addition, we observed that Npnt was localized in the acellular primary stroma suggesting that prior to its expression by the migratory pNC in the cornea, the optic cup and lens vesicle are the primary sources of Npnt in the nascent corneal ECM. The link between the primary stroma and the presumptive corneal epithelium and endothelium was established in classical studies, which showed that the lens and corneal endothelium induce the corneal epithelium to synthesize collagen and glycosaminoglycans into the underlaying space (Fitch et al., 1994; E. Hay et al., 1979; E. D. Hay & Revel, 1969; Toole & Trelstad, 1971). Our expression analyses suggest that the primary stroma sequesters Npnt initially secreted by the optic cup and the lens, and exposes it to pNC during corneal development.

Itgα8 is expressed in the spinal cord, optic nerve, retina, urogenital and digestive systems, and in the head epidermis (Bossy et al., 1991; Ogawa et al., 2018). α8β1 is known to modulate epithelial-mesenchymal interactions during kidney and pharyngeal development (Müller et al., 1997; Talbot et al., 2016) and to promote migration of mesangial and smooth muscle cells (Bieritz et al., 2003; Zargham & Thibault, 2006). We report for the first time that *Itgα8* is expressed by pNC during ocular development. Since Itgα8 heterodimerizes with Itgβ1 subunit (Bossy et al., 1991), which is robustly expressed in the periocular mesenchyme and migratory pNC (Doane & Birk, 1994), we infer that α8β1 signaling functions during corneal development in response to Npnt secreted in the primary stroma. Following delamination from the neural tube, cranial neural crest migrate in response to Fn and laminin (Duband & Thiery, 1982; Newgreen & Thiery, 1980; Sternberg & Kimber, 1986) via integrin receptors such as α1β1, α4β1, α5β1, αVβ1 (Alfandari et al., 2003; Delannet et al., 1994; Desban & Duband, 1997; Kil et al., 1996; Lallier et al., 1994). Although neural crest cells are initially highly migratory, they remain relatively stationary upon their localization in the periocular region. Given that *Itgα8* expression coincides with the onset of pNC migration, our results indicate its potential role in their ingression into the developing cornea.

### Npnt promotes pNC migration via RGD domain and Itgα8

The three major factors that contribute to corneal thickness during development are pNC migration, cell proliferation, and synthesis of the secondary stroma by the keratocytes. We showed that disruption of *Npnt* expression in the anterior ocular tissues via RCAS-mediated knockdown resulted in decreased corneal thickness. In addition to functioning in epithelial-mesenchymal interactions, Npnt has been shown to promote migration in various tissues including vascular endothelial cells during osteogenesis (Kuek et al., 2016), infiltration of immune cells into the liver (Hong et al., 2020; Inagaki et al., 2013), and cancer metastasis (Magnussen et al., 2020; Mei et al., 2020; Wang et al., 2018). Given that Npnt is localized in the primary stroma during corneal development, we hypothesized that it may play a potential role in pNC migration. In agreement with our hypothesis, our *in vitro* migration assays confirmed that robust pNC migration from mesenchyme explant occurred on Npnt substrate but was abrogated in the presence of the Itgα8 inhibitor. We also observed that knockdown of *Itgα8* reduced pNC migration into the cornea and phenocopied the reduction of corneal thickness observed following *Npnt* knockdown. In addition, we found that reduction in corneal thickness was accompanied by decrease in cell count, but cell density and proliferation were not affected, suggesting that Npnt/Itgα8 signaling mediates pNC migration during corneal development. Given that the primary function of keratocytes is to synthesize the corneal ECM comprised of collagens and proteoglycans, and represents approximately 90% of the corneal thickness (Fini, 1999; Funderburgh et al., 2003; Kao, 2010), our observation that there was no change in stromal cell density raises a possibility that matrix synthesis by the pNC that differentiated into keratocytes is not a major contributing factor to the reduction in corneal thickness.

Furthermore, our overexpression studies showed the opposite effect whereby the full-length construct caused increased corneal thickness at E9. We also found that increased corneal thickness persisted at E15 when Npnt appears to be downregulated in the cornea. Surprisingly, the E7 corneas were not affected in the overexpression studies. This could be caused by limited pNC migration potential due to their expression of Neuropilin1, which prevents them from entering the corneal environment that contains a repulsive Semaphorin3A signal (Lwigale & Bronner-Fraser, 2009). As observed in the knockdown studies, there was no differences in cell proliferation between control and *Npnt^oe^* corneas, suggesting that there was an increase of pNC migration. The mutant Npnt-RAE construct overexpresses approximately the same size protein as the endogenous Npnt, while maintaining its MAM domain to support ECM-ECM interactions, but it did not impact cornea thickness. This indicates that expression of high levels of protein does not affect cornea thickness, further implicating that Npnt functions in directing cell migration specifically through its RGD sequence during early cornea development. The reduction in cell count and density could be attributed to the mutant RAE domain outcompeting the expression of the endogenous RGD by the pNC. Similarly, the truncated version of Npnt containing only the EGF-like repeats did not affect cornea thickness, further conforming that Npnt signals through the RGD domain and Itgα8 to modulate pNC migration into the cornea.

Precise coordination of multiple signals from surrounding ocular tissues orchestrate the spatiotemporal migration and differentiation of multipotent pNC during the formation of the avian corneal endothelium and stromal keratocytes. Disruptions in the sequence of these early events can lead to significant defects in corneal development. Our study provides the first characterization of Npnt/Itgα8 function in pNC that contribute to the cornea. Despite being surrounded by an ECM rich in Fn, pNC remain relatively immobile in the periocular region until they are competent to respond to Npnt cues via the expression of α8β1. This transformation may be crucial for the timely induction of directional migration of pNC towards the gradient of additional cues generated by Npnt in the primary stroma (Fig.7A, B), and therefore plays a vital role in segregating the cornea progenitors from the rest of the periocular mesenchyme. Our results also show that Npnt subsequently localizes to the basement membrane of the corneal epithelium. Future studies will investigate the potential role of Npnt in epithelial-mesenchymal interactions between the corneal epithelium and stroma, potentially as a result of signaling through the EGF-like domain to EGFR receptors in the epithelial cells.

## MATERIALS AND METHODS

### Chick embryos

All experiments were performed using fertilized White Leghorn chicken eggs (*Gallus gallus domesticus*) obtained from Texas A& M Poultry Center (College Station, TX). Eggs were incubated at 38 °C in humidified conditions until the desired stages. Animal studies were approved by the Institutional Animal Care and Use Committee (IACUC) at Rice University.

For Npnt and Itgα8 knockdown and overexpression studies, eggs were incubated for approximately 24-26hrs to obtain 3-somite stage or HH8 (Hamburger & Hamilton, 1951), then windowed as previously described (Spurlin & Lwigale, 2013). A few drops of Ringer’s solution containing 100 U/mL penicillin and 100μg/mL streptomycin (PenStrep, ThermoFisher Scientific) were added to embryos to maintain hydration. Embryos were injected with viral constructs (see below) using a Picospritzer III pneumatic microinjection system (Parker Hannifin) in the space between the vitelline membrane and cranial region, ensuring that the neural tube and adjacent ectoderm were completely covered (Fig. 2A). Injected embryos were re-incubated and collected for GFP screening at the desired stages of development ranging between embryonic day (E)5-E15. Only the corneas showing robust GFP expression were used for subsequent analyses.

### Histology, Hematoxylin and Eosin staining, and Immunohistochemistry

Embryos were collected at desired stages and eyes were dissected in Ringer’s saline solution. Samples collected for Hematoxylin and Eosin (H&E) staining were fixed overnight at 4 °C in modified Carnoy’s fixative (60% ethanol, 30% formaldehyde, and 10% glacial acetic acid). Afterwards, the tissues were dehydrated in ethanol series, cleared in Histosol (National Diagnostics), embedded in paraffin blocks, and sectioned at 10 μm thickness. Sections were stained with Hematoxylin for 15 seconds, counterstained with Eosin for 40 seconds, dehydrated in ethanol series, and mounted with Cytoseal (ThermoFisher Scientific) for imaging. For Npnt immunohistochemistry, eyes were fixed in methanol-acetic acid (MAA) fixative (98% methanol, 2% glacial acetic acid) that was chilled on dry ice. Samples were maintained in MAA fixative at - 80 °C for at least two days, then were gradually warmed to room temperature before further dehydration in ethanol series and embedding in paraffin blocks. Samples were sectioned at 10 μm, rehydrated, then immunostained with Npnt antibody (1:100; orb221700, Biorbyt) following standard procedures. For all the other immunohistochemistry and phalloidin staining, samples were fixed in 4% PFA overnight at 4 °C, embedded in paraffin, and sectioned as described above. Immunostaining with anti-GFP (1:500; A-6455, Invitrogen), and anti-fibronectin (1:30; B3/D6, Developmental Studies Hybridoma Bank) was used to detect protein expression. Phalloidin-568 (1:200; A-12380, Invitrogen) and DAPI (Roche) was used to show total cell distribution in wholemount and sectioned tissue, respectively. The following secondary antibodies (Invitrogen) were used at 1:200: Alexa-488 goat anti-rabbit IgG, Alexa-594 goat anti-rabbit IgG, Alexa-594 goat anti-mouse IgG2a, and Alexa-488 goat anti-mouse IgG1.

### Section *In situ* Hybridization

Eyes were fixed in modified Carnoy’s fixative, embedded in paraffin, and sectioned as described above. Riboprobes were generated from gene fragments using cDNA pooled from E7 anterior eyes and cloned into TOPO-PCRII (Invitrogen). Digoxigenin (DIG)-labeled riboprobes were synthesized following manufacturer’s protocol (DIG Labeling Kit, Roche). Primers used to generate the riboprobes are listed in Supplementary Table 1. Sections were hybridized with riboprobe at 52 °C (*Npnt*) and 60 °C (*Itgα8*) overnight. Hybridization was detected using anti-DIG antibody conjugated with alkaline phosphatase (Roche) and color was developed with 5-bromo-4-chloro-3-indolyl phosphate/nitro blue tetrazolium (BCIP/NBT; Sigma). Following color development, sections were fixed with 4% PFA, mounted in Cytoseal and imaged using an Axiocam mounted on an AxioImager2 microscope (Carl Zeiss).

### Production of RCAS virus

Replication-Competent ASLV long terminal repeat with a Splice acceptor (RCAS) (Hughes et al., 1987) virus was used for stable and prolonged expression of short hairpin (sh)RNA constructs used for knockdown and for the overexpression studies. To produce RCAS viral stocks, chick fibroblasts (DF-1 cells; Lot 58217603, ATCC) were cultured in Dulbecco’s modified eagle medium (DMEM) supplemented with 10% fetal bovine serum (FBS, Invitrogen), 100 U/mL penicillin and 100 μg/mL streptomycin (referred to here as complete DMEM, ThermoFisher Scientific). Cells were transfected at about 70% confluency with plasmid DNA containing the RCAS constructs using Lipofectamine 3000 (Invitrogen). Transfected cells were grown for four days to enable viral replication. Media containing viral particles were collected on subsequent days, pooled, and centrifuged at 21,000 rpm (Beckman) for 1.5 hour at 4 °C to concentrate the virus. Virus pellet was resuspended in DMEM and stocks at approximately 1-7×10^6^ Ifu/ml were stored in −80 °C until use.

### Generation of shRNA and viral constructs

shRNA target sequences used for *Npnt^kd^* and *Itgα8^kd^* (Supplementary Table 2), were designed using the BLOCK-iT RNAi Designer tool (ThermoFisher Scientific) and linked to their reverse complementary sequence by a loop sequence, TTCAAGAGA. A short termination sequence and restriction enzyme sites were added at each end to create fragments that were ligated into a pSLAX shuttle vector (Hughes et al., 1987) and to add a chick U6 (Cu6) promoter and a GFP reporter. The shRNA sequences were cloned into the RCAS vector (Supplementary Fig. 2A,B) by homologous recombination using a CloneEZ kit (GenScript) as previously described (Kwiatkowski et al., 2017; Ojeda et al., 2017). The *Npnt^oe^* overexpression constructs were either generated by substitution of the shRNA with a full-length coding sequence of Npnt in the vector above or using a modified vector where the full-length Npnt protein was driven by a promoter within the viral long terminal repeats (LTRs) with eGFP linked by an IRES motif (Supplementary Fig. 2A, B). Constructs containing either mutated Npnt with the RGD sequence changed to RAE (PR<u>GD</u>VFIPRQPGVSNNLFEIL<u>E</u>I<u>E</u>R to PR<u>AE</u>VFIPRQPGVSNNLFEIL<u>A</u>I<u>A</u>R) or the truncated version of Npnt containing only the N-terminal EGF-like repeats domain (Npnt-EGF) were commercially synthesized (Genscript) and cloned into the modified RCAS vector. All constructs were validated using primers for Npnt and Itgα8 (Supplementary Table 3).

### BrdU staining

Bromodeoxyuridine (BrdU) was used to assay for cell proliferation. The BrdU solution was prepared at a final concentration of 10 μM in complete media. Eyes were collected at desired stages and injected into the anterior chamber between the cornea and lens with BrdU solution (approximately 50-100 μl depending on development stage), then cultured in BrdU solution for 2 hours at 37 °C. Eyes were rinsed in phosphate buffered solution (PBS), fixed in 4% PFA, embedded in paraffin, and sectioned as described above. BrdU-positive cells were identified by immunohistochemistry using anti-BrdU antibody (1:30; G3G4, Developmental Studies Hybridoma Bank). Sections were counterstained with DAPI.

### *In vitro* explant culture

Embryos were collected at E4 in Ringer’s solution and the anterior eyes were dissected and digested in dispase (1.5mg/mL, Worthington) for 10 minutes at 37 °C. The periocular mesenchyme was isolated by physically removing the presumptive cornea epithelium, lens, and the optic cup. The mesenchyme ring was further trimmed using tungsten needles to obtain cells that are proximal to the presumptive cornea, then dissected into approximately 120×120 μm explants. Nunc Lab-tek II 8-well chamber slides (Sigma) were coated with 1.5 μg/cm^2^ with poly-D lysine (MP Biomedicals) for 1 hour at room temperature, followed by recombinant mouse or human Npnt (R&D Systems) at 1.5 μg/cm^2^ for 2 hours at 37 °C. The α8β1 peptide inhibitor following the 23-mer sequence P**RGD**VFIPRQPTN**DLFEIFEIER** (Sato et al., 2009) was commercially generated (Genscript) and used at a 10 μM working concentration. Mesenchyme explants were transferred to the coated slides containing complete media, with and without α8β1 inhibitor and incubated at 37 °C in a humidified tissue culture incubator with 5% CO_2_ for 12 hours. Explants with migratory cells were imaged with a Rebel T6s camera (Canon) mounted on an Axiovert 40C microscope (Carl Ziess). The magnitude of cell migration from explants was calculated by measuring the density of cells in a 175 μm x 175 μm area located 270 μm from the center of the explant.

### Time-lapse video microscopy

Mesenchyme explants were prepared and cultured as described above in media containing 0.1 μg/ml Hoechst dye solution (ThermoFisher Scientific) used to label all nuclei. Chamber slides containing attached explants were imaged at 3 minutes and 27 second intervals for 17 hours using an FV1200 laser scanning microscope (Olympus) with a stage top incubator (Tokai Hit). Movies were generated using IMARIS Software (Oxford Instruments).

### Quantification of Corneal Measurements

#### Corneal thickness

Differences in corneal thickness between control and knockdown samples was determined by averaging measurements taken at three separate locations along the cornea. All measurements were taken perpendicular to the radial curvature of the cornea. Since corneal thickness is not always uniform in overexpression samples, measurements were taken at three locations across the center of the thickened region.

#### Cornea cell counts and density

Nuclei labeled by the DAPI staining of corneal sections were used to count stromal cells within a width of 200 μm along the entire height of that region. Cells were either counted manually or using the threshold particle analysis in ImageJ Software in which nuclei from the epithelium and endothelium are discounted from the final tally. Cell density was determined by dividing the number of nuclei and the area, which was measured by multiplying the value of the corneal thickness by the width of the selected region. Comparison of cell counts and density between the anterior vs posterior corneal region was determined by taking similar areas for all corneas, which was set at a value of 20% the average thickness of control corneas.

#### Cell proliferation

Cell proliferation was determined in corneal sections by first utilizing the cell count technique described above. The total number of cells was represented by all the DAPI-positive nuclei, out of which the BrdU-positive cells was quantified. Cell proliferation was calculated as a percentage of total the DAPI-positive nuclei that were BrdU positive.

### Software and Statistics

ImageJ Software was used to measure the staining intensity or tissue morphological features, such as cornea thickness and cell density. The number of cornea sections analyzed are summarized in each figure legend. All statistical analyses were conducted using GraphPad Software. Data are presented as scatterplot with mean values. Statistical significance was determined by two-tailed unpaired Student’s t-test was used to compare differences between means. Samples with p-values < 0.05 were considered significant.

## ACKNOWLEDGEMENTS

We would like to thank members of Lwigale lab for the helpful discussions and suggestions on this project. We would also like to thank the Warmflash lab for use of the FV1200 laser scanning microscope for live imaging of periocular mesenchyme explants. This work was funded by National Institutes of Health Grants R01 EY031381 and EY022158 (to P.Y.L).

## COMPETING INTERESTS

No competing interests to declare.

---

**Movie 1: Migration of pNC from mesenchyme explant on Npnt-coated substrate.** Time-lapse movie was taken over 17 hours with images taken every 3 minutes and 27 seconds. The cells are shown in bright-field and fluorescent Hoechst nuclear staining (blue). Relates to Fig. 3E.

**Movie 2: Migration of pNC from mesenchyme explant on Npnt-coated substrate in the presence of α8β1 inhibitor.** Time-lapse movie was taken over 17 hours with images taken every 3 minutes and 27 seconds. The cells are shown in bright-field and fluorescent Hoechst nuclear staining (blue). Relates to Fig. 3F.

**Source Data 1: Contains statistical analysis reported in Figures 2-6.**

**Source Data 2: Contains gel images for data reported in Supplementary Figure S2C.**

## SUPPLEMENTARY MATERIAL

**Supplementary Table 1.**
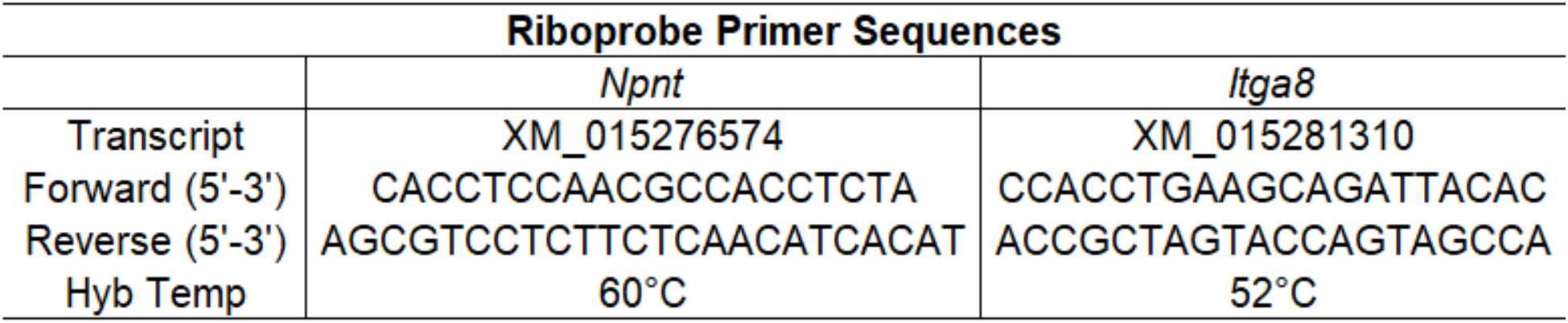
Primer sequences for riboprobe synthesis.

**Supplementary Table 2.**
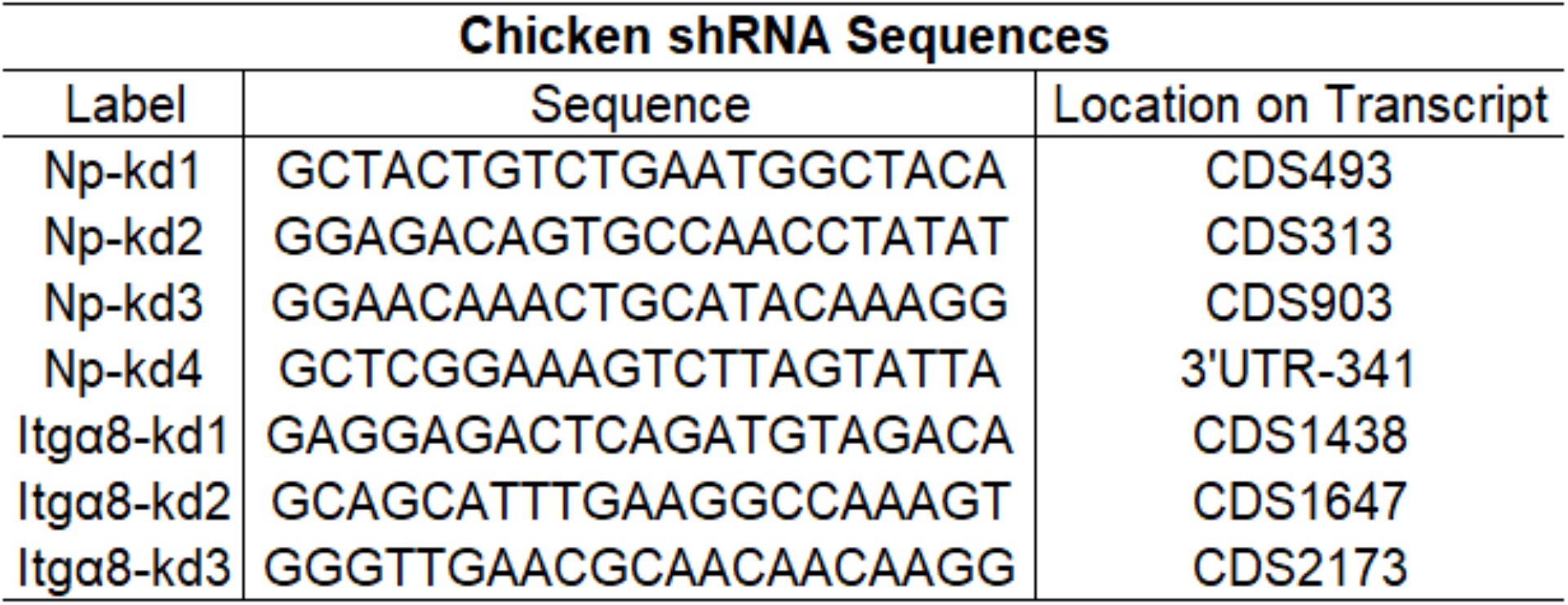
shRNA target sequences used for knockdown studies.

**Supplementary Table 3.**
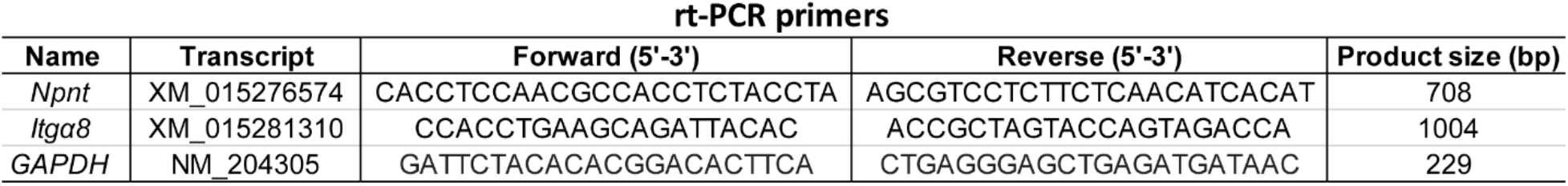
Primers used to validate *Npnt* and *Itgα8* knockdown efficiency. rt-PCR primers.

**Supplementary Figure S1.**
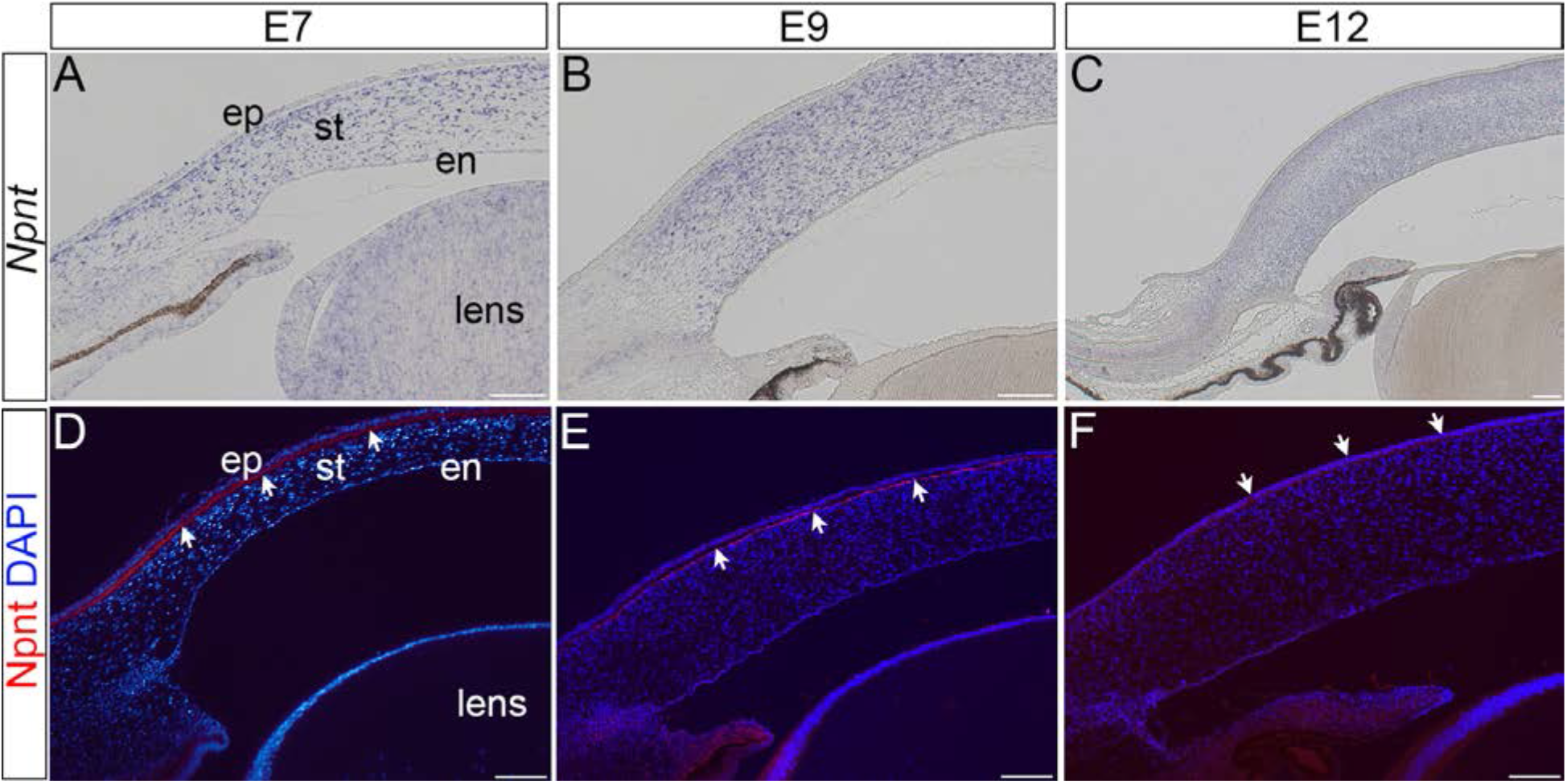
Expression of Npnt transcripts and protein during late stages of development of the chick cornea. **(A-C)** Section in situ hybridization showing localization of Npnt in the stroma of E7, E9, and E12 corneas. **(D-F)** Immunohistochemistry showing that Npnt (red) is localized in the ECM directly adjacent to the corneal epithelium at E7 and E9 (**D** and **E**, arrows), and in the corneal epithelium at E9 (F, arrows). Sections are counterstained with DAPI (blue). Scale bars represent 100 μm. Abbreviations: ep, epithelium; st, stroma; en, corneal endothelium.

**Supplementary Figure S2.**
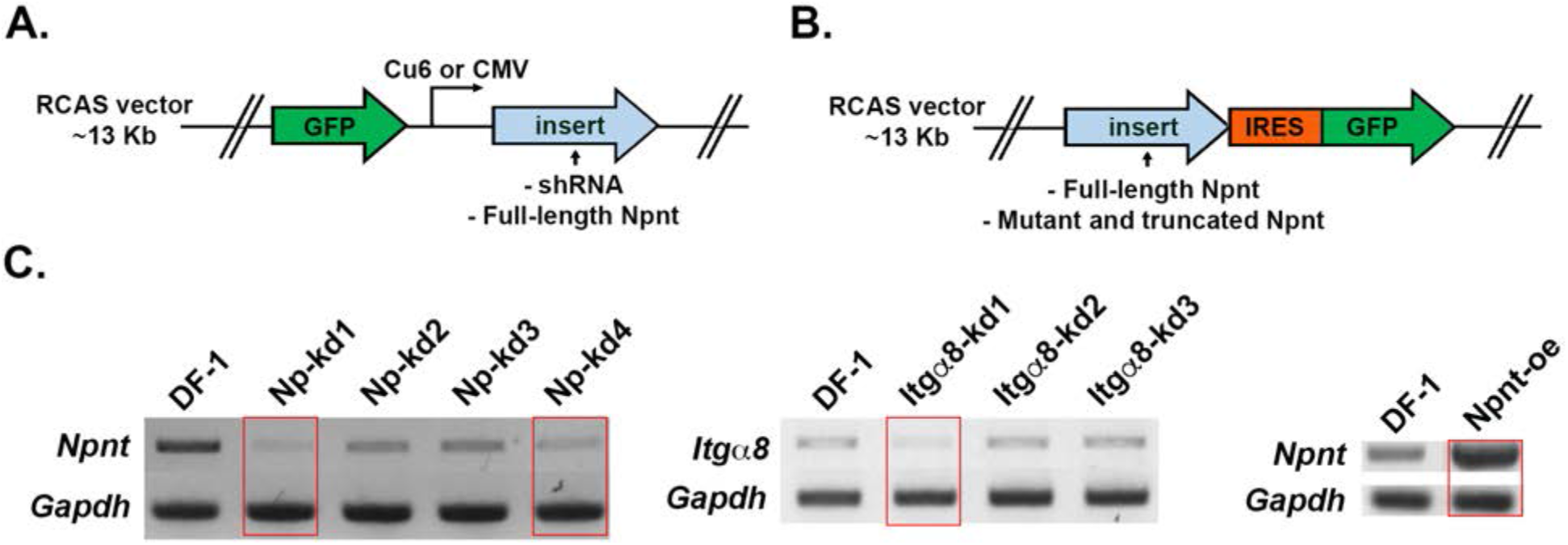
Diagram of RCAS vectors used for expression of either: **(A)** GFP alone (control), or GFP together with inserts of shRNA or full-length Npnt. **(B)** The full-length or mutated and truncated versions of Npnt driven by the viral LTR promoter and GFP driven by the IRES promoter. **(C)** For RTPCR, cells plated in 30 mm dishes were homogenized in 1 mL TRIzol and RNA was isolated following manufacture’s protocol. RNA samples were treated with Turbo DNA-*free* kit (Invitrogen) to remove residual genomic DNA and cDNA pools were generated using SuperScript First Strand System (Invitrogen). Semi-quantitative PCR was conducted using HotStart-IT Taq polymerase (Affymetrix). Knockdown efficiency was measured on 2% agarose gel using GAPDH (glyceralde-hyde-3-phosphate dehydrogenase) as a loading control. Red rectangles indicate the constructs that were chosen for the knockdown and overexpression experiments. Primers for RT-PCR are located in Table 1.

**Supplementary Figure S3.**
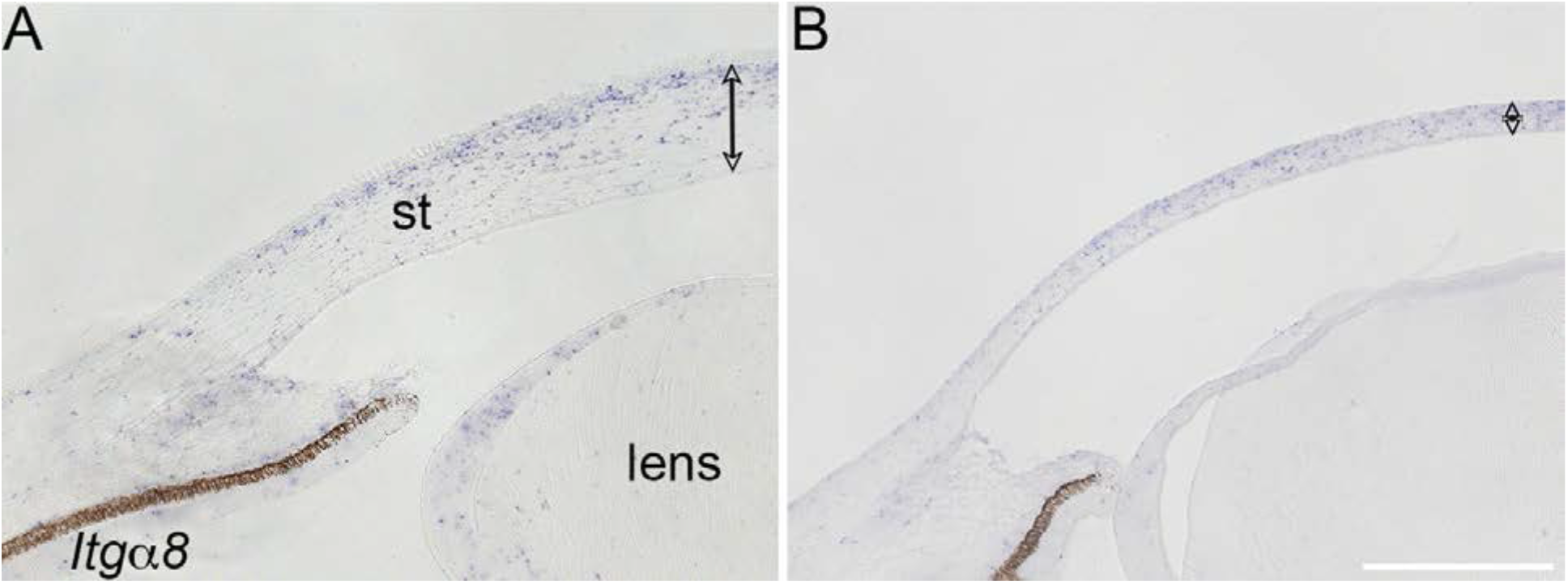
Validation of *Itgα8* knockdown in vivo. Stage 8 chick embryos were injected in the cranial region with control and *Itgα8*-shRNA viral constructs, and re-incubated until E7. Section in situ hybridization was performed using riboprobes for *Itgα8*. (A) Section of control cornea showing *Itgα8* expression in the corneal stroma, iris and lens epithelium. (B) section of *Itgα8^kd^* cornea showing reduced expression of *Itgα8* in the thin cornea, iris and lens epithelium. The double sided arrows indicate cornea thickness. Abbreviation: st, corneal stroma. Scale bar, 200 μm.

## Notes

### Competing Interest Statement

The authors have declared no competing interest.

### Summary of Updates

To add supplementary files

