## Supplementary material for "Nephronectin-Integrin α8 signaling is required for proper migration of periocular neural crest cells during chick corneal development": Source data 2

### Supplementary Figure S2C Source Data

#### Npnt-kd

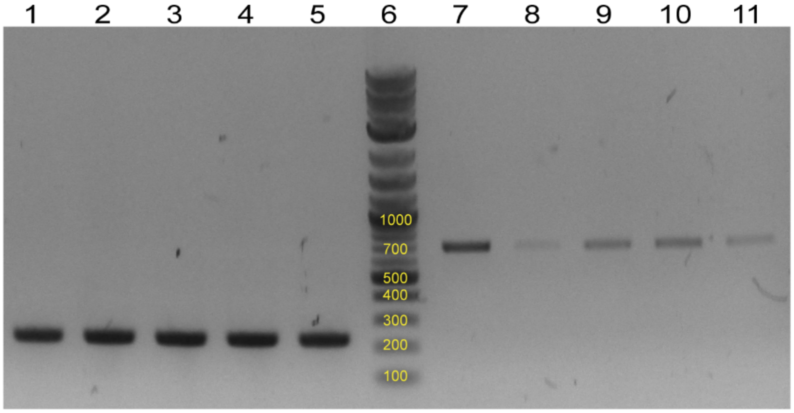


Lanes 1-5: GAPDH (229bp); lane 6: NEB 1kb plus ladder; lanes 7-11: Npnt (708bp)

cDNA sample:
Lanes 1, 7: DF-1
Lanes 2, 8: Npnt-kd1
Lanes 3, 9: Npnt-kd2
Lanes 4, 10: Npnt-kd3
Lanes 5, 11: Npnt-kd4

#### Itgα8-kd

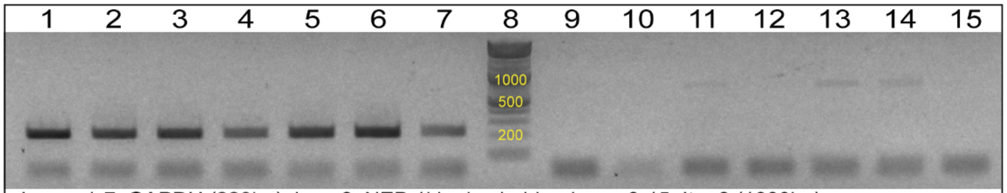


Lanes 1-7: GAPDH (229bp); lane 8: NEB 1kb plus ladder; lanes 9-15: Itgα8 (1000bp)

cDNA sample:
Lanes 3, 11: DF-1
Lanes 4, 12: Itgα8-kd1
Lanes 5, 13: Itgα8-kd2
Lanes 6, 14: Itgα8-kd3

*Samples not used in this manuscript: 1, 2, 7, 9, 10, 15*

#### Npnt-OE

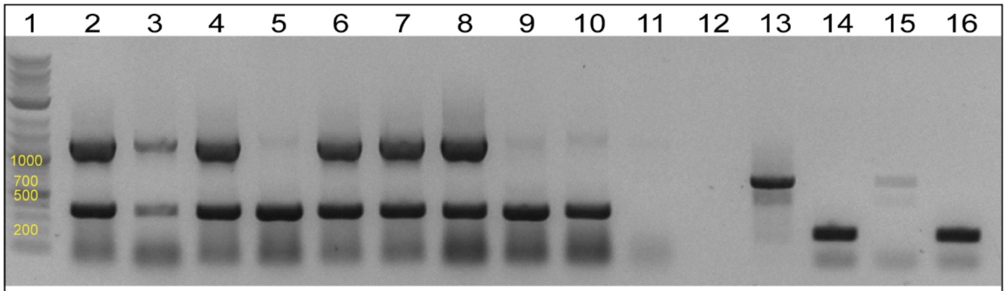


Lane 1: NEB 1kb plus ladder; lanes 2-12: not used in this manuscript; lanes 13, 15: Npnt (708bp); lanes 14, 16: GAPDH (229bp)

cDNA sample:
Lanes 13, 14: Npnt-OE
Lanes 15, 16: DF-1
